# One-shot analysis of translated mammalian lncRNAs with AHARIBO

**DOI:** 10.1101/2020.04.20.050062

**Authors:** Luca Minati, Claudia Firrito, Alessia Del Piano, Alberto Peretti, Simone Sidoli, Daniele Peroni, Romina Belli, Francesco Gandolfi, Alessandro Romanel, Paola Bernabò, Jacopo Zasso, Alessandro Quattrone, Graziano Guella, Fabio Lauria, Gabriella Viero, Massimiliano Clamer

## Abstract

A vast portion of the mammalian genome is transcribed as long non-coding RNAs (lncRNAs) acting in the cytoplasm with largely unknown functions. Surprisingly, lncRNAs have been shown to interact with ribosomes, encode uncharacterized proteins, or act as ribosome sponges. These functions still remain mostly undetected and understudied owing to the lack of efficient tools for genome-wide simultaneous identification of ribosome-associated lncRNAs and peptide-producing lncRNAs.

Here we present AHARIBO, a method for the detection of lncRNAs either untranslated, but associated with ribosomes, or encoding small peptides. Using AHARIBO in mouse embryonic stem cells during neuronal differentiation, we isolated ribosome-protected RNA fragments, translated RNAs and corresponding *de novo* synthesized polypeptides. Besides identifying mRNAs under active translation and associated ribosomes, we found and distinguished lncRNAs acting as ribosome sponges or encoding micropeptides, laying the ground for a better functional understanding of hundreds lncRNAs.

## INTRODUCTION

An incredibly small fraction of the mammalian genome is protein-coding (< 3 %), while the number of potentially functional non-coding genes remains unclear (Djebali et al., 2012). LncRNAs are defined as non-coding RNA exceeding 200 nt. They have gained much attention because of their role in a variety of cellular processes, from chromatin architecture (Minajigi et al., 2015) to mRNA turnover (Kleaveland et al., 2018) and translation (Ingolia et al., 2011). Typically, lncRNAs are abundant transcripts (Iyer et al., 2015) that display short and not evolutionarily conserved ORFs with minimal homology to known protein domains (Guttman and Rinn, 2012). The majority of the lncRNAs are localized in the cytoplasm (Carlevaro-Fita et al., 2016), where they are supposed to be untranslated. Ribosome profiling (RIBO-seq), which provides positional information of ribosomes along transcripts (Clamer et al., 2018; Ingolia et al., 2012a), identified several ribosome-associated lncRNAs (Bazzini et al., 2014; Ingolia et al., 2011; Lee et al., 2012; Zeng et al., 2018). A handful of lncRNAs have been shown to be involved in the modulation of translation (Carrieri et al., 2012; Yoon et al., 2012), while others have been found to be potentially translated (Anderson et al., 2015; Aspden et al., 2014; Bazin et al., 2017; van Heesch et al., 2019; Ingolia et al., 2011; Nelson et al., 2016; Ruiz-Orera et al., 2014). As coding RNAs, lncRNAs can be associated with fully translating of with translationally silent ribosomes (Chandrasekaran et al., 2019; Chen et al., 2020; Jiao and Meyerowitz, 2010; Kapur et al., 2017). Hence, the potential involvement of lncRNAs in translation, increases the complexity of the mammalian translatome and proteome. Unfortunately, classical RIBO-seq approaches cannot distinguish between lncRNAs producing peptides from those that are sequestering ribosomes (lncRNA bound to ribosomes without translation), thus acting as ribosome sponges. Complementary proteomics approaches, such as mass spectrometry, can help defining and quantitatively monitoring the production of peptides, but are less sensitive techniques than RNA sequencing (van Heesch et al., 2019; Slavoff et al., 2013). Therefore, proteomics and RIBO-seq alone cannot unravel the wide functional range of cytoplasmic lncRNAs associated with the translation machinery.

To fill this gap, we developed AHARIBO (<u>AHA</u>-mediated <u>RIBO</u>some isolation), a combination of protocols to simultaneously isolate RNAs and nascent proteins associated with translationally active ribosomes. AHARIBO is based on the isolation of ribosomes trapped with their nascent peptides, by means of the incorporation of the non-canonical amino acid L-azidohomoalanine (AHA), followed by parallel RNA-seq, ribosome profiling and proteomics.

We applied AHARIBO to human and mouse cells and showed that it enables to: i) purify translating ribosomes via nascent peptide chains, ii) analyze co-purified RNAs and proteins for transcriptome / *de novo* proteome-associated studies, and iii) detect the regulatory network of lncRNAs translated or associated with ribosomes.

## RESULTS

### Nascent peptide labelling and separation of the ribosome complex with AHARIBO-rC

To simultaneously purify ribosomes under active translation, associated RNAs and corresponding growing peptide chains, we optimized a protocol in HeLa cells (Fig 1A). Briefly, the protocol consists in the following phases: i) incubation with a methionine-depleted medium, ii) addition of the methionine analogue AHA, iii) on-ribosome anchorage of nascent peptide chains by means of a small molecule, iv) cell lysis and AHA “copper-free click reaction” (Jewett and Bertozzi, 2010) for v) final ribosome capture with magnetic beads. We reasoned that the protocol for isolating ribosomes through AHA can be used to obtain information from nascent peptides, constitutive components of the ribosomes, mRNAs or lncRNAs associated with ribosomes. For this reason, we optimized several parameters along the entire protocol, from washing steps to nuclease treatments (Fig. 1A), to isolate (1) the full translational complex (ribosomes, ribosome-associated proteins, nascent peptides and RNAs; called AHARIBO-rC, <u>R</u>ibosomal <u>C</u>omplex), (2) the *de novo* synthesized proteome (called AHARIBO-nP, <u>N</u>ascent <u>P</u>roteins) and (3) ribosome-protected fragments (RPFs) (called AHARIBO RIBO-seq, <u>RIBO</u>some profiling by <u>seq</u>uencing).

**Figure 1.**
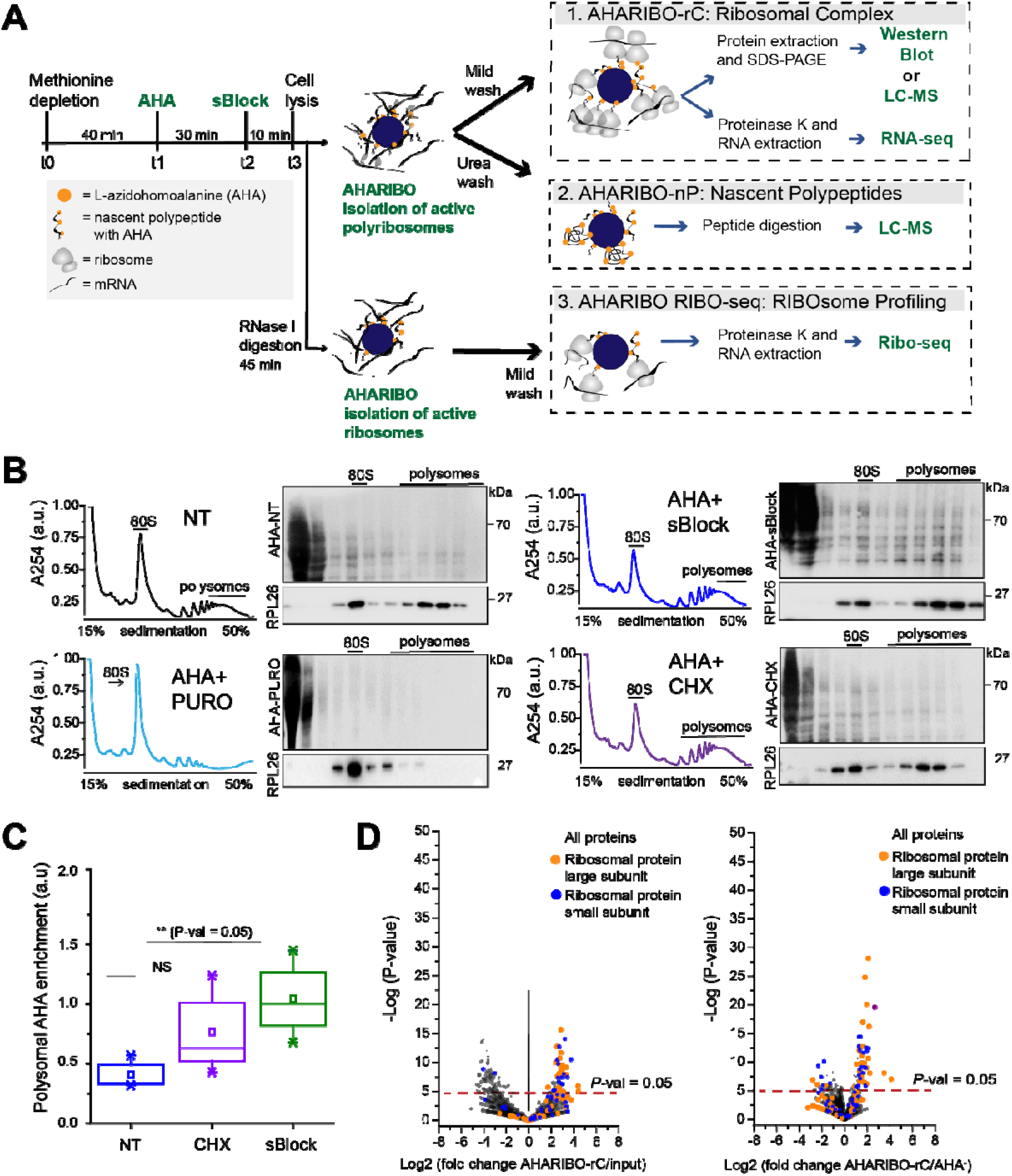
AHA labeling of nascent peptide chains and ribosome separation. A) Schematic representation of the AHARIBO experimental workflow. After methionine depletion, AHA incubation and sBlock treatment, cell lysates can be processed for (1) AHARIBO-rC: isolation of translational complex (ribosomes, ribosome-associated proteins, nascent peptides and RNAs); (2) AHARIBO-nP: isolation of *de novo* synthesized proteins; (3) AHARIBO RIBO-seq. B) Sucrose gradient profiles resulting from absorbance measurement at 254 nm. On the right of each profile, SDS-PAGE of protein extracts from each fraction of the profile. After ultracentrifugation on a 15-50% sucrose gradient, HeLa cell lysates were fractionated (1 mL fraction) and the total AHA protein content of each individual fraction was visualized after SDS-PAGE. Staining of the membrane was performed by biotin cycloaddition followed by streptavidin-HRP. RPL26 protein was used as ribosome marker in the different fractions. C) Box plot showing the AHA signal enrichment in the polysomal region of the profiles obtained from cells treated with either CHX or sBlock or no drugs (NT). Results are shown as a median (± SE) of 3 independent replicates. D) Volcano plots showing the -Log (P-value) versus the relative abundance of AHARIBO-rC-isolated proteins. Data are compared with input (starting AHA-containing lysate, left) or with the non-specific signal from streptavidin-coated beads incubated with cell lysate not supplemented with AHA (AHA).

To minimize the amount of AHA-tagged proteins already released from ribosomes and achieve optimal on-ribosome polypeptide stabilization, we tested different times of AHA incubation and compared the effect of two small molecules (namely cycloheximide (CHX) and sBlock, an anisomycin-based reagent), which are known to inhibit the activity eukaryotic ribosome, while keeping polypeptides bound to the translating ribosomes (Garreau de Loubresse et al., 2014; Grollman, 1967; Seedhom et al., 2016)

We observed that 30 min is the optimal incubation time for sufficient AHA incorporation and maximum RNA recovery (Supplementary Fig. 1A-C). Then, we compared the drug efficiency of stabilization of the nascent peptide by co-sedimentation profiling of AHA-tagged polypeptides with ribosomes (Fig 1B). As a control, cells were treated in parallel with puromycin to cause ribosome disassembly and release of the growing peptide chains (Fig. 1B) (Blobel and Sabatini, 1971). According to literature, we found that both CHX and sBlock are able to stabilize AHA-peptides on ribosomes and polysomes (i.e. actively translating complexes) (Biever et al., 2020; Mathias et al., 1964). The efficiency to anchor polypeptides on ribosomes in CHX and sBlock treated cells was 50% higher compared to untreated cells, confirming that the drugs effectively stabilize nascent polypeptides (Fig. 1C). The high signal observed in light fractions is caused by AHA-labeled proteins released from ribosomes. As expected, in puromycin-treated samples the AHA signal was mainly detected in the first two fractions, proving that the observed signal in the heavier fractions of CHX- and sBlock-treated cells is not caused by a diffusion of AHA-labeled peptides. Since sBlock outperformed CHX in anchoring efficiency (Fig. 1C), we used sBlock in all further experiments.

Prompted by the evidence that it is possible to stably anchor nascent peptides on ribosomes by means of a small molecule, we validated the ability of AHARIBO-rC to isolate RNAs and proteins associated with the translation complex. To this aim, we performed a label-free liquid chromatography-mass spectrometry (LC-MS) analysis on captured proteins. Enrichment analysis of AHARIBO-rC proteins relative to the input and to the non-specific signal (AHA) showed that ribosomal proteins belonging to both the large and the small ribosome subunits are indeed more abundant in AHARIBO-rC samples than the controls (Fig. 1D). Gene ontology (GO) analysis of these proteins revealed that terms related to translation (biological process), nucleic acid binding (cellular function) and ribonucleoprotein complex (cellular component) are enriched in AHARIBO-rC compared to the control (AHA), confirming efficient pulldown of proteins involved in translation (Supplementary Tab. 1). LC-MS results were also confirmed by Western blot analysis on proteins involved in translation (RPS6, RPL26 and eEF2) (Supplementary Fig. 1D).

Then, we investigated the possibility to use AHARIBO-rC to determine the translational status of cultured cells. To address this question, we down-regulated protein synthesis by treating HeLa cells with arsenite (Ar), which induces translational inhibition and stress granules formation (Wang et al., 2016). Consistent with our previous evidence, we observed a reduction of RNA captured in Ar-treated cells relative to the control (Supplementary Fig. 1E-F). In line with this finding, qRT-PCR analysis showed about 50% reduction in 18S rRNA levels when translation is inhibited (i.e. in Ar AHARIBO-rC pulldown samples compare to not arsenite treated cells) (Supplementary Fig. 1G).

### AHARIBO-nP: genome-wide portrayal of the *de novo* synthesized proteome

Motivated by the evidence that AHARIBO-rC can be used to isolate *bona-fide* translationally active ribosomes, we further tested the method genome-wide on mouse embryonic stem cells (mESCs) under a basal condition and mESCs differentiated into early neurons (EN) (Tebaldi et al., 2018) (Supplementary Fig. 2A). We analyzed both AHARIBO-rC isolated RNA and newly synthesized polypeptides associated with actively translating ribosomes by RNA-seq and LC-MS, respectively. The protocol for the isolation of the *de novo* synthesised polypeptides (named AHARIBO-nP) is based on urea washing to remove all proteins that are not nascent polypeptides (Supplementary Fig. 2B). In parallel, we isolated and analyzed the global translatome as the RNA extracted after 30% sucrose cushioning of cytoplasmatic lysates (Wang et al., 2013) and the global proteome, measured by pulsed SILAC (pSILAC) (Schwanhäusser et al., 2009) (Fig. 2A and 2B).

**Figure 2.**
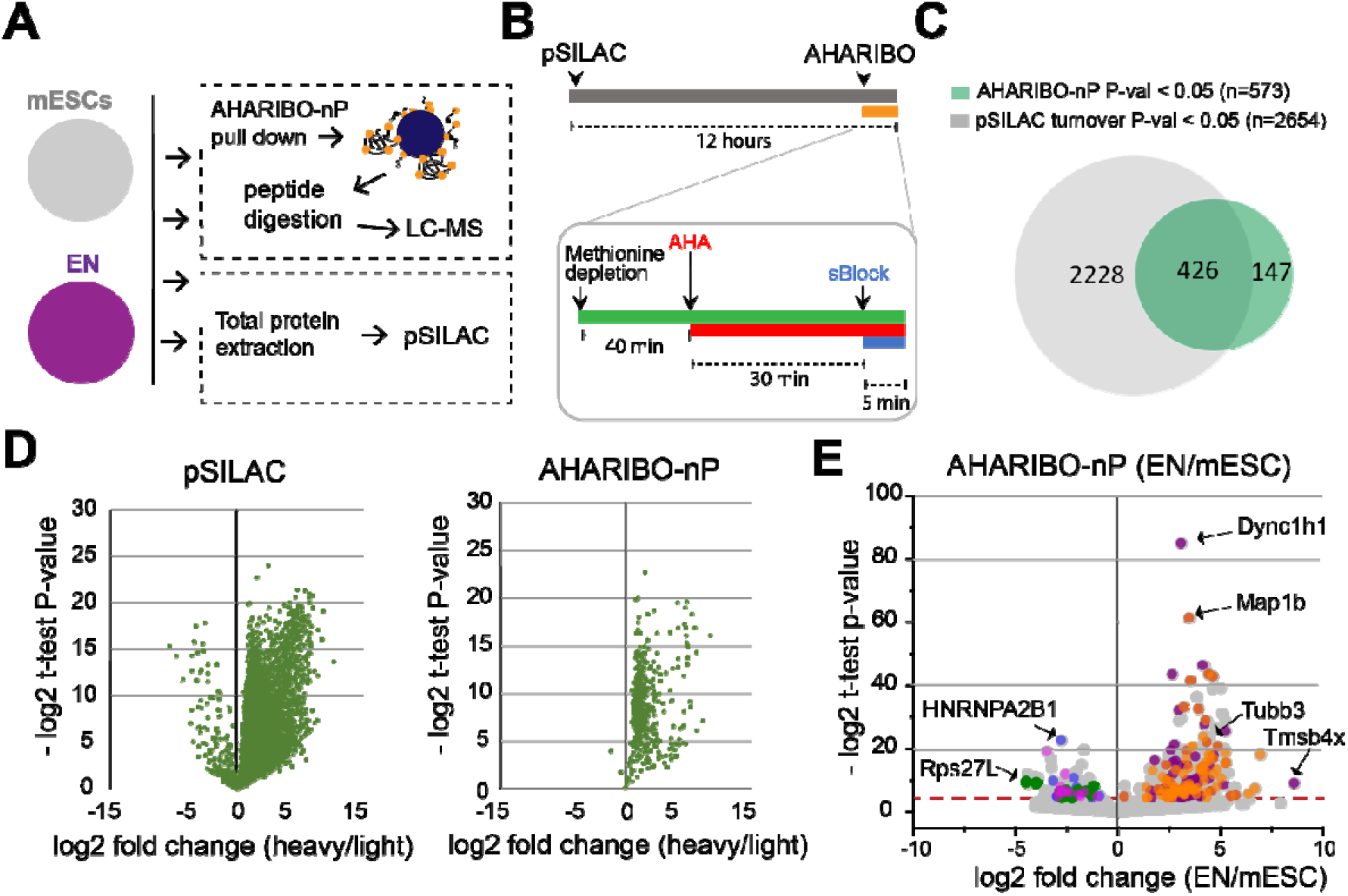
AHARIBO-nP and pSILAC approaches. A) Schematic representation of the experimental workflow. B) Schematic representation of combined cell treatments for pSILAC and AHARIBO-nP. C) Venn diagram representing the number of proteins identified by AHARIBO-nP and pSILAC. D) Volcano plots displaying for each protein the -log2 p-value against the fold changes of protein turnover (heavy/light) in pSILAC proteome (left) and AHARIBO-nP (right) for mESCs. E) Volcano plot displaying for each differentially expressed protein (EN/mESC) the AHARIBO-nP proteome versus -log2(p-value). Red broken line indicates P-value < 0.05. Orange and purple dots represent up-regulated proteins involved in cytoskeleton organization (GO:0007010) and neurogenesis (GO: 0022008) respectively. Blue, green and magenta dots represent down-regulated proteins related to RNA processing (GO:0006396), protein synthesis (GO:0006412) and mouse pluripotency (WP1763) respectively. Grey dots, all proteins. Gene Ontology (GO) enrichment analyses were performed using PANTHER and Enrichr.

We identified 2417 newly synthesized proteins (heavy/light, P-val < 0.05) within the 24 hours of pSILAC treatment. As expected, differentiated cells (EN) showed a reduced turnover (Supplementary Fig. 2C). Then, we performed an enrichment analysis of ENs versus mESCs (EN/mESC) and identified 2667 and 573 differentially expressed proteins in pSILAC and AHARIBO-nP samples, respectively (Fig. 2C). The lower number of proteins identified with AHARIBO-nP compared to pSILAC is most probably related to the shorter time of incubation with AHA (30 min) compared to pSILAC (24 hours) and is consistent with previous observations from similar pulldown enrichment strategies (Bagert et al., 2014; Rothenberg et al., 2018). Of note, 74 % of the AHARIBO-nP identified proteins is in common with the pSILAC dataset (Fig. 2C). Accordingly, we observed high abundance of heavy amino acids in AHARIBO-nP (Fig. 2D) and a significantly higher heavy/light ratio in AHARIBO-nP compared to pSILAC (Supplementary Fig. 2D), suggesting that AHARIBO-nP is indeed able to capture the *de novo* synthesized proteome. We found positive fold changes for proteins captured by AHARIBO-nP during differentiation (Fig. 2E). Interestingly, among these proteins, several are known to be expressed during the early stages of development of the nervous system (e.g. Map1b, Tubb3 and, Dync1h1). The GO analysis of differentially expressed proteins showed that proteins involved in cytoskeleton organization and neurogenesis were upregulated (Fig. 2E), further confirming the reliability of AHARIBO-nP in monitoring *de novo* protein expression.

Collectively, these results show that *de novo* synthesized proteins isolated using AHARIBO-nP portray the phenotypic changes induced by cellular differentiation, and that AHARIBO-nP protocol is suitable to monitor dynamic changes in protein expression by LC-MS analysis.

### Combination of AHARIBO-rC and AHARIBO-nP: parallel genome-wide analysis of translated RNAs and *de novo* synthesized proteome

Prompted by previous results, we tested if mRNAs purified using AHARIBO-rC can be a good proxy of protein levels. To this aim, we compared AHARIBO-rC RNA and the global translatome with AHARIBO-nP in mESCs during differentiation.

To exclude any bias related to protein length, we checked whether AHARIBO-nP preferentially captures either long or short proteins. We analyzed peptide size against the AHARIBO-rC RNA enrichment versus the global translatome (Fig. 3A), a value reflecting the extent to which AHARIBO-rC RNA differs from the standard analysis. Positive values were associated with long proteins (> 200 aa) in both mESCs and ENs. A 3.5 % of the proteins showed length of less than 100 aa. Proteins shorter than 100 aa account for approximately 10% of the total mouse proteome, with an average protein size in eukaryotes of about 300 aa (Frith et al., 2006). Since in all eukaryotes proteins are initiated with a methionine residue, virtually any protein can be captured as soon as the nascent peptide exits the ribosome (when it reaches a length of about 35-40 aa). In about 70 % of the proteome, the N-terminal methionine is co-translationally cleaved when the peptide is at least 50 aa long by the enzyme methionine aminopeptidase (Wild et al., 2020), while the remaining 30 % retains the methionine (Martinez et al., 2008). Therefore, there is a good probability for at least one AHA residue to be available for each peptide when the blocking drug is added to the cell medium, enabling the capture of the polypeptide outside the ribosome exit tunnel. Our results confirm the effectiveness of the method and the efficiency in capturing polypeptides without major biases.

**Figure 3.**
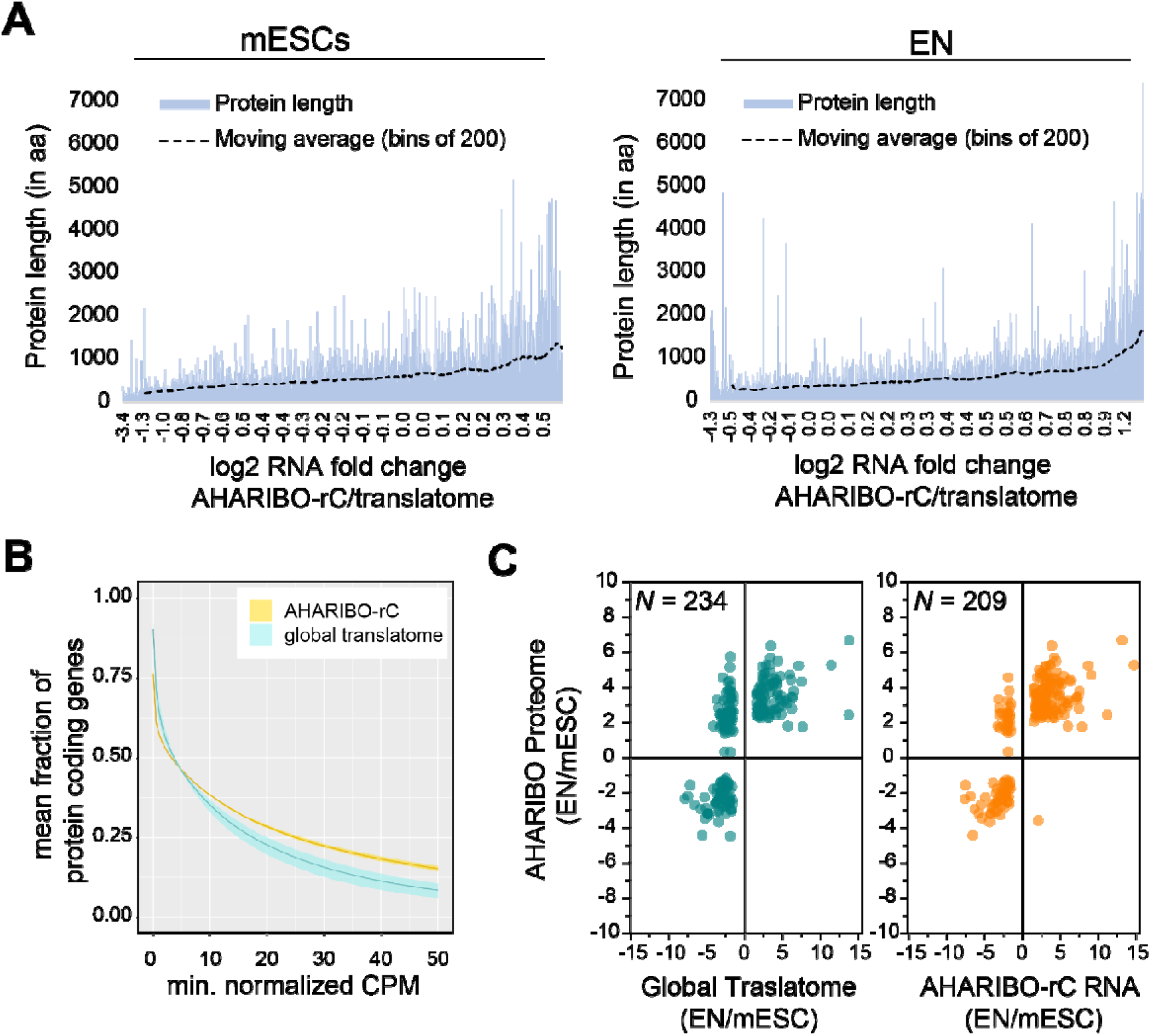
AHARIBO-rC RNA vs *de novo* proteome analysis. A) Protein length against increasing fold changes of AHARIBO-rC RNA/global translatome for mESCs (left) and ENs (right). B) Fraction of remaining protein-coding genes at increasing CPM thresholds in ENs. The mean of n = 3 (translatome) and n = 3 (AHARIBO-rCAHARIBO-rC) biological replicates is reported. Data are represented as mean ± SD. (shadow). C) Scatter plot of RNA fold change (global translatome on the left, AHARIBO-rC on the right) compared to protein fold change (AHARIBO-nP). N, number of DEGs with P-val < 0.05.

To further exclude any bias related to RNA abundance, we measured how efficiently AHARIBO-rC captures coding transcripts compared to the global translatome. We analysed the two datasets using increasing abundance thresholds in EN, and observed that AHARIBO-rC is comparable to the global translatome for low abundant transcripts and it can better capture more abundant transcripts (Fig 3B). In undifferentiated mESCs AHARIBO-rC performs as the global translatome (Supplementary Fig. 3A), confirming the efficiency of successful isolation of protein-coding RNAs genome-wide.

Finally, we tested whether the RNA isolated with AHARIBO-rC can predict the *de novo* synthesized proteome. After comparing differentially expressed genes (DEGs, with P-Val < 0.05) obtained by AHARIBO-rC RNA during mESC differentiation to the AHARIBO-nP proteome, we observed that AHARIBO-rC RNA is a good proxy of the newly synthetized proteome (Pearson’s correlation *r =* 0.75) (Fig. 3C and Supplementary Fig. 3B). In particular, we found that AHARIBO-rC RNA presents less uncoupled genes (up-RNA and down-protein or down-RNA and up-protein) than the global translatome (Fig 3C and Supplementary Fig. 3C), thus recapitulating protein changes faithfully.

### Combined AHARIBO approaches define the functional role of lncRNAs in translation

LncRNAs are known to be in some cases associated with ribosomes. Ribosome-associate lncRNAs play a role in the regulation of translation (Ingolia et al., 2011; Pircher et al., 2014; Zeng et al., 2018) or in the production of short peptides (Chen et al., 2020; van Heesch et al., 2019). Based on our evidence that a combination of AHARIBO approaches can simultaneously detect RNAs under active translation and peptides in the process of being produced, we applied our methods to detect ribosome-associated and translated lncRNAs.

From the analysis of lncRNAs isolated by AHARIBO-rC, we identified about 400 differentially expressed (DE) lncRNAs during neuronal differentiation (Supplementary Fig. 4A). Among the top-5 DE lncRNAs, we found *Pantr1* and *Lhx1os*, known to be involved in neuronal development (Biscarini et al., 2018; Carelli et al., 2019).

**Figure 4.**
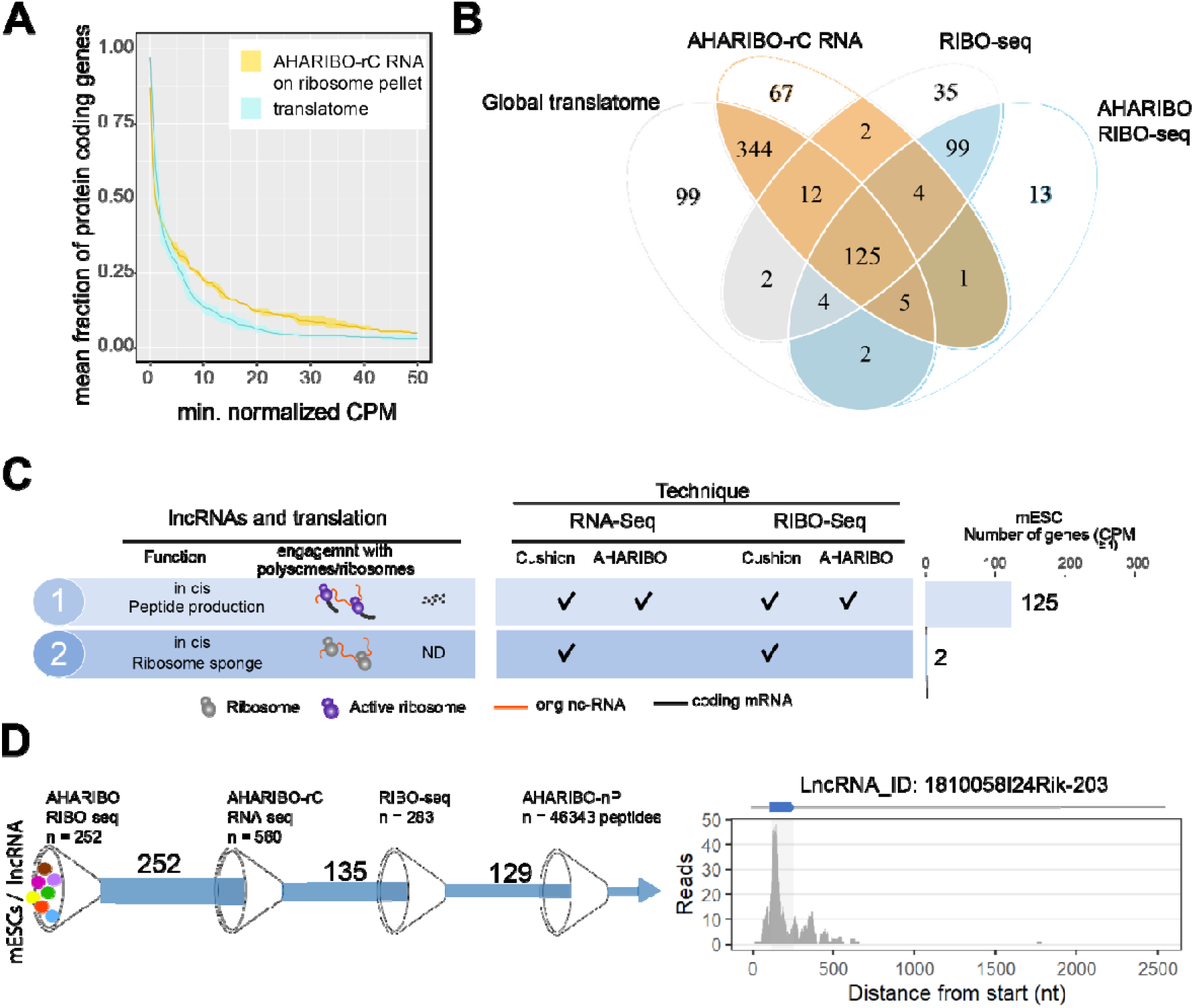
The AHARIBO platform can be used to detect ribosome-interacting lncRNAs. (A) Fraction of remaining non-coding genes at increasing CPM thresholds in EN. AHARIBO-rC was performed on the ribosome pellet after cushioning. The mean of n = 3 independent replicates is reported. Data are represented as mean + SD. (shadow) B) Venn diagram with the number of lncRNAs (with at least 1 CPM) identified in each category (RNA-seq, AHARIBO-rC, RIBO-seq, AHARIBO RIBO-seq). C) Left, Classification of lncRNAs interacting with ribosomes and relative detection through the different AHARIBO and standard approaches. D) Left, schematic representation of the number of mESC lncRNAs in common between AHARIBO RIBO-seq, AHARIBO-rC RNA, standard RIBO-seq, and finally validated by LC-MS. Right, example a AHARIBO RIBO-seq trace of the IncRNA 1810058124Rik displaying the hallmarks of translation including reads distribution along the entire transcript and the accumulation of reads in the known short open reading frame (shadow area and blue arrow on top).

To identify potentially translated lncRNAs, we applied the abundance threshold analysis to the subset of AHARIBO-rC non-coding RNAs in common with a published dataset (n = 270) of lncRNA with ribosome footprints (RPFs), identified by ribosome profiling in mESCs (Ingolia et al., 2011) (Supplementary Fig. 4B). The analysis of the 100 lncRNAs in common between the two datasets showed a stronger enrichment of AHARIBO-rC lncRNA ribosome footprints in the AHARIBO-rC than in the global translatome (Fig. 4A and Supplementary Figure 4C), meaning that only a subset of lncRNAs are potentially translated. This result suggest that a fraction of non-coding transcripts efficiently isolated with AHARIBO-rC are potentially translated.

To understand if and how lncRNAs interact with ribosomes, we performed RIBO-seq after AHARIBO pulldown (named AHARIBO RIBO-seq), and standard RNA-seq and RIBO-seq (on inputs) in mESCs. With AHARIBO RIBO-seq samples we identified an absolute number of 252 lncRNAs within RPFs. For protein-coding genes, both standard and AHARIBO RIBO-seq enriched for RPFs. The two datasets correlates well (R^2^ = 0.96) and the P-site density showed the expected 3nt periodicity in the coding sequence, thus confirming the capability of AHARIBO protocol in capturing ribosomes (Supplementary Figure 4D). By intersecting AHARIBO RIBO-seq data with those obtained from standard methods (RIBO-seq, RNA-seq after sucrose cushioning) and AHARIBO-rC, we identified 125 common putative translated lncRNAs (Fig. 4B). The vast majority of these lncRNAs have no already known function, demonstrating the our approach could be useful to unravel new and translation-related lncRNAs. Two lncRNAs (9330151L19Rik and Gm9776) were identified only by standard RIBO-seq and RNA-seq but not with AHARIBO (Fig. 4C), suggesting that these are not translated, but rather loaded with idle ribosomes.

To validate the coding potential of lncRNAs which are in common between AHARIBO and standard RIBO-seq (n = 228) (Fig. 4D), we first *in silico* translated the transcripts in all frames to find potential ORFs with a canonical start codon (AUG). Translated sequences were then semi-trypsin digested *in silico,* screened to remove contaminants and then manually annotated to find confident matching spectra from the AHRIBO-nP protein dataset. MS-based proteomics detected five non-canonical peptides with corresponding ribosome footprints (Gm42743, Gm26518, B230354K17Rik, D030068K23Rik, 1810058124Rik), out of the about 46,000 collected spectra. The low number of validated transcript could be due to challenges in detecting the trypsin-digested products from short, non-canonical coding sequences. From the short list of 129 in common with all AHARIBO protocols and standard RIBO-seq (Fig. 4D), we identified at a high degree of confidence a known micropeptide of 47 aa (Fig. 4D) derived from a lncRNA expressed in murine macrophages. This micropeptide was recently detected and characterized as a relevant protein in the modulation of the innate immunity in mice (Bhatta et al., 2020).

Altogether, our data confirm that the three diverse and complementary AHARIBO approaches represent a unique method to identify ribosome-associated and translated lncRNAs.

## Discussion

LncRNAs can be localized in the nucleus or in the cytoplasm. When localizing in the nucleus they can modulate transcription, pre-mRNA splicing or act as scaffold for protein interaction during chromatin organization (Sun et al., 2018). When localizing in the cytoplasm, the majority is associated to polysomes (Carlevaro-Fita et al., 2016) where lncRNA can be used to produce or not to produce proteins (Chen et al., 2020; Ingolia et al., 2011). Numerous lncRNAs are misannotated as non-coding but contain short ORFs encoding for micropeptides with biological relevance in cancer (D’Lima et al., 2017; Huang et al., 2017), bones development (Galindo et al., 2007), macrophage activity (van Solingen et al., 2018), metabolism (Magny et al., 2013; Nelson et al., 2016) and DNA repair (Slavoff et al., 2014). Different methodological approaches have been developed to quantify the variations of mRNA abundance (e.g., RNA-seq, RNA imaging) (Amit Blumberg et. al, 2019; Jao and Salic, 2008; Morisaki et al., 2016; Wu et al., 2016), RNA engagement with the translational machinery (e.g., ribosomal profiling, polysomal profiling) (Arava et al., 2003; Clamer et al., 2018; Eden et al., 2011; Taniguchi et al., 2010), and protein synthesis (e.g., pSILAC, PUNCH-P, BONCAT) (Aviner et al., 2013; Dieterich et al., 2006; Schwanhäusser et al., 2009; Yan et al., 2016). Nevertheless, available technologies hardly capture in a single experimental setup the dynamics of translation across different biological conditions, the translation of unannotated coding transcripts and translation-related functions of lncRNAs. Now that is widely accepted that a portion of the genome annotated as non-coding (> 97 %), can result in a complex transcriptome partially engaged with ribosomes (Chen et al., 2020; Djebali et al., 2012; Iyer et al., 2015), exosome sequencing and ribosome profiling should include micropeptide detection.

With AHARIBO, we introduce a strategy for the selective isolation of active ribosomes via nascent peptide chain for more comprehensive interrogation of lncRNA biology and proteogenomic studies. Our data show that AHARIBO serves as a flexible tool to detect translated RNAs, to identify lncRNAs bound to elongating ribosomes and to detect *de novo* synthesized proteins in a fast and accessible way. The intersection of standard RIBO-seq, RNA-seq and AHARIBO approaches allowed us to identify translated lncRNA as well as few example of lncRNA sequestering ribosomes without evidence of translation. Although RNA sequencing analysis is not supported by the same diversity of LC-MS data, we successfully identify a mouse-specific micropeptide reported to originate form a IncRNA ORF as a case study, confirming the effectiveness of AHARIBO.

Overall, we provide evidence that AHARIBO is a comprehensive and reliable toolkit suitable for downstream orthogonal approaches (RNA-seq, RIBO-seq and LC-MS), empowering scientists to shed light onto the functional complexity of the mammalian genome and lncRNA translation.

## SUPPLEMENTARY INFORMATION

**Figure S1.**
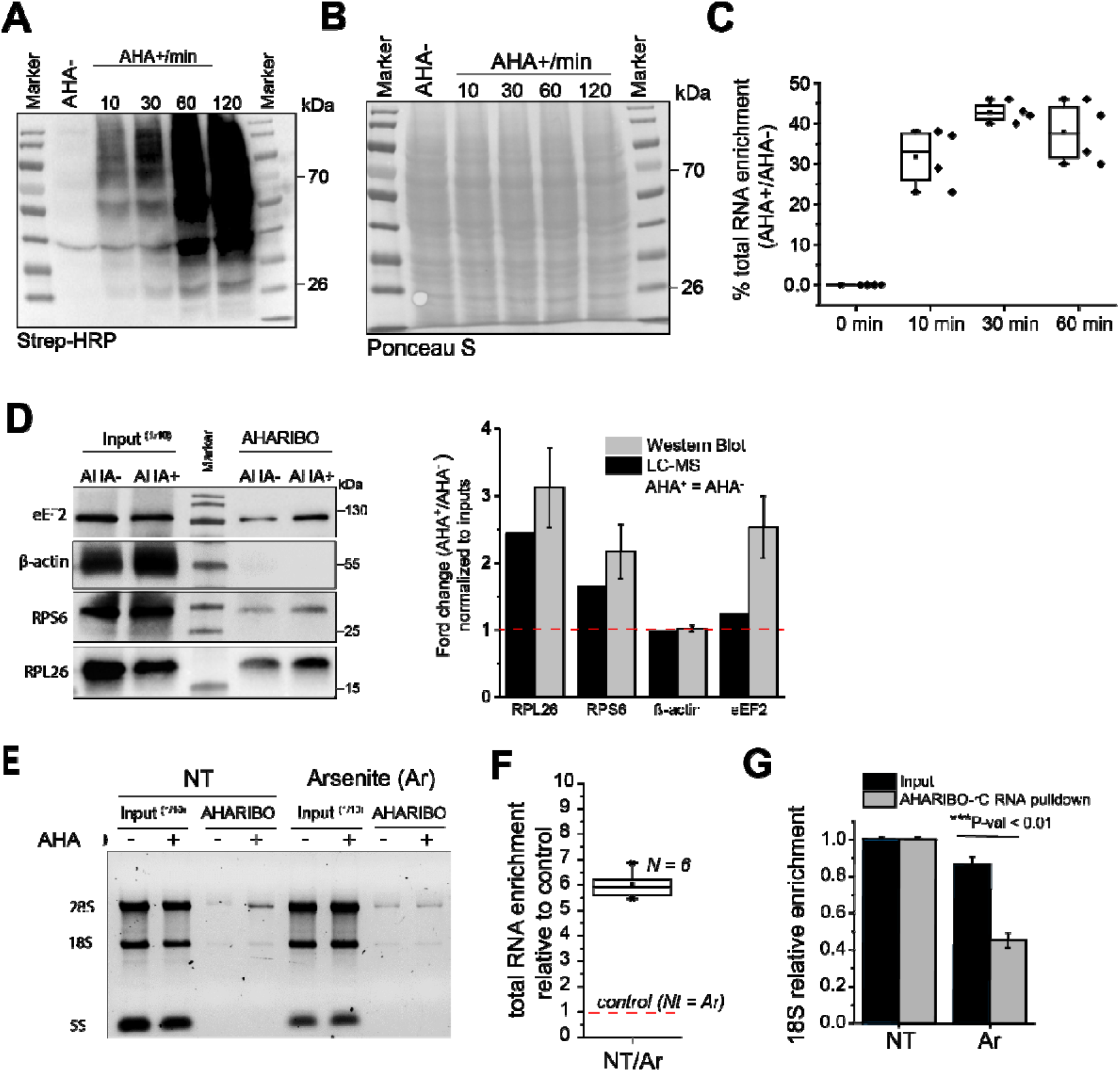
AHA incorporation, validation of AHA and RNA capture. A) Labeling of nascent peptides in cells treated with AHA (250 μM) at different incubation times (10, 30, 60, 120 min). After SDS-PAGE of cell extracts, AHA residues were biotinylated by on-membrane cycloaddition based “click chemistry” and detected by streptavidin-HRP. B) Ponceaus S staining of the membrane reported in A. C) RNA enrichment in AHATIBO-tC pulldown at different AHA incubation times (10, 30, 60 min) compared to control (AHA^*^) and reported as % of input (1/10 of total RNA). D) Immunoblotting analysis of proteins captured by AHARIBO-rC and obtained from cells either treated or not treated with AHA. A lysate volume corresponding to 1/10th of the volume processed for pulldown was used as reference inputs. On the right, histogram reporting the relative abundance of RPL26, eEF2, beta-actin and RPS6 in LC-MS analysis and quantitative analysis of Western Blot data (n = 3; error bars, ±SD). E) Agarose gel electrophoresis of total RNA extracted from input lysates (1/10 of the total lysate volume) and lysates subjected to AHARIBO pulldown, obtained from cells either treated or not treated with AHA. NT, non-treated cells; Ar, arsenite-treated cells. F) Total RNA enrichment after AHARIBO-rC pulldown of lysates obtained from unstimulated cells over cells treated with arsenite. For each condition, cells were either treated or not treated with AHA. Signal ratios (AHA^+^/AHA^−^) for each pulldown sample were normalized to the respective inputs. NT, non-treated. Square box, mean; stars, 1-99% percentile. G) 18S rRNA qRT-PCR analysis of RNA extracted from lysates subjected to AHARIBO-rC pulldown and input lysates, obtained from unstimulated cells or cells subjected to arsenite treatment. For each condition, cells were either treated or not treated with AHA. For each sample, 18S AHA^+^/AHA^−^ signal ratios were normalized to the input and to the housekeeping gene HPRT1. NT, Non-treated; Ar, arsenite.

**Figure S2.**
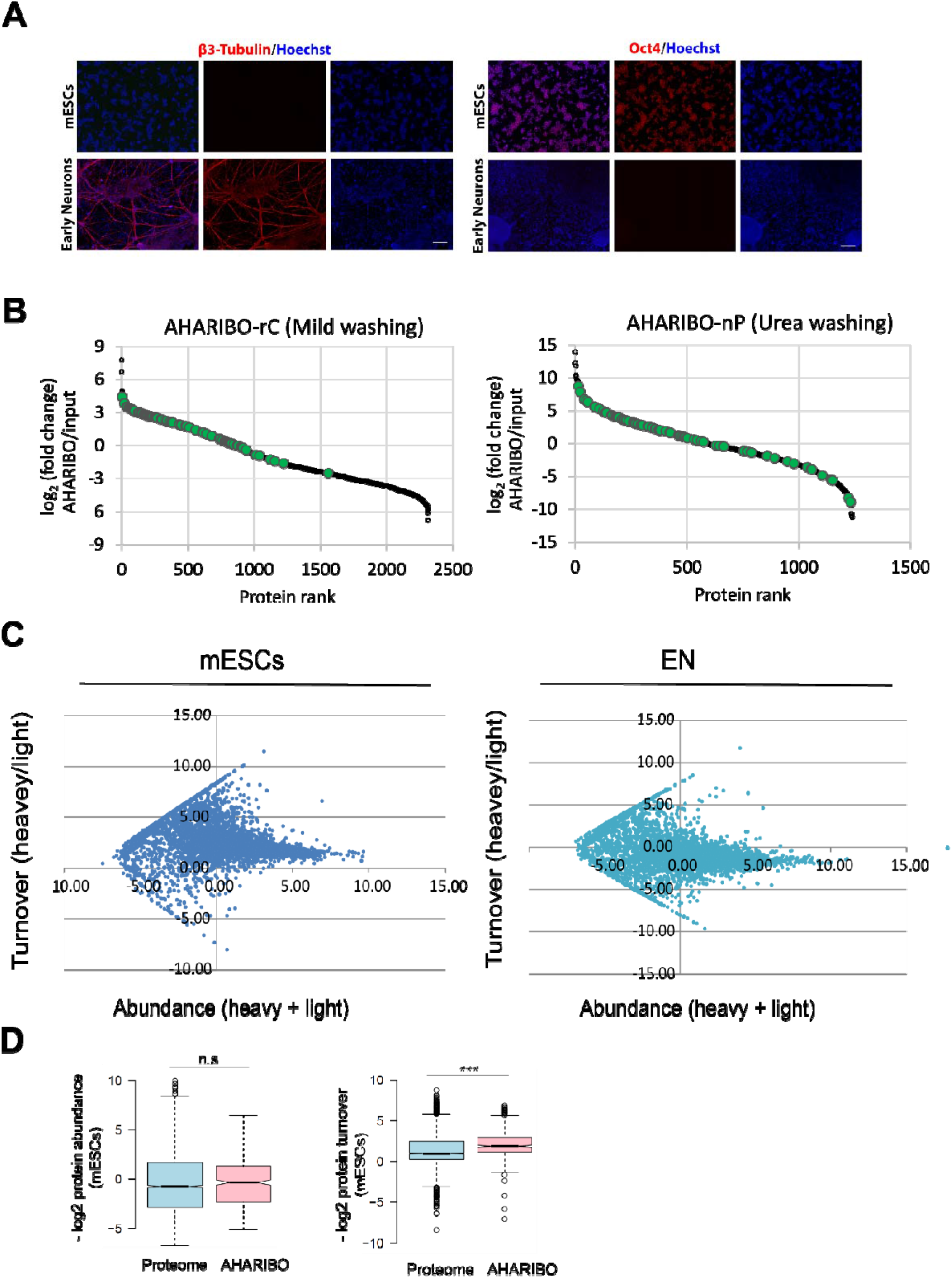
Proteomic analysis. A) Immunofluorescence for mESC (Oct4) and neuronal (ß3-tu bu I in) marker expression on self-renewing mESCs and 15DIV mESC-derived neurons. Scale bar: 200 μm. B) Rank plot of fold change of full proteome (black dots) and ribosomal proteins (green dots), comparing AHARIBO pulldown versus input samples, mild washing (left) and urea washing (right). Since AHARIBO-rC LC-MS analysis might cause an underestimation of the *de novo* synthesized proteome due to the enrichment of abundant ribosomal proteins, newly synthetized proteins bound to DBCO-conjugate magnetic beads were separated from ribosome subunits by harsh washing conditions (8M urea) before tryptic digestion and LC-MS analysis. The effectiveness of the washing procedure was confirmed, since no evident enrichment of ribosomal proteins in the pull-down was observed. C) The scatter plots represent protein abundance versus protein turnover in mESCs (left) and ENs (right). D). Normalized protein abundance (left) and turnover distribution (right) as determined by pSILAC and AHARIBO. ***P-Val < 0.001.

**Figure S3.**
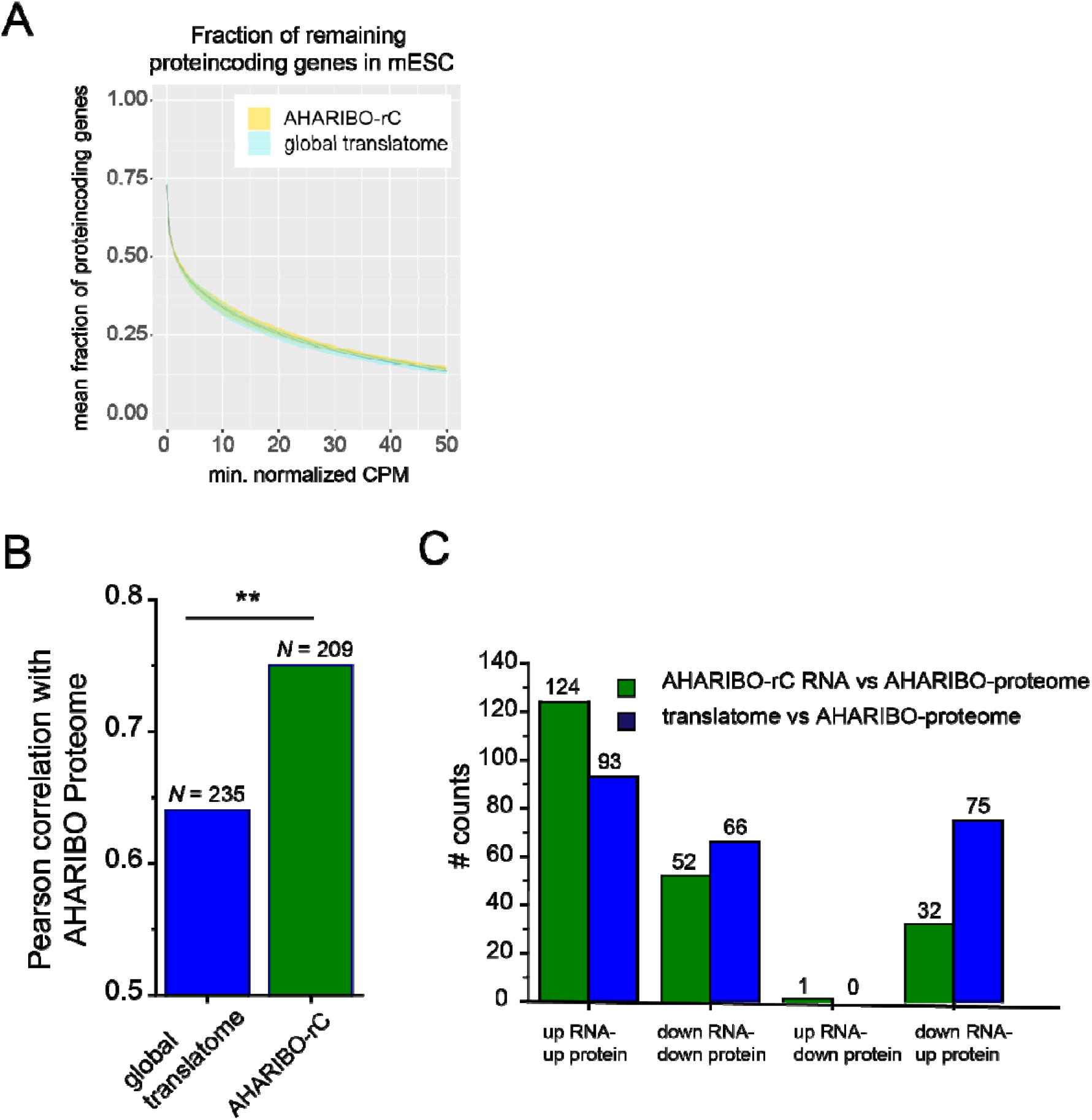
RNA-seq and protein coding RNA analysis. A) Fraction of remaining protein coding genes at increasing CPM thresholds in mESCs. The mean of n = 3 biological replicates is reported. Shadows: data are represented as mean ± SD. B) Histogram showing Pearson’s correlation analysis of AHARIBO-nP protein fold change (EN/mESC) determined by mass spectrometry versus global translatome and AHARIBO-rC RNA fold change (EN/mESC) determined by RNA-seq. N, number of DEGs. P-val < 0.05. C) Histogram of the numberof DEGs (EN/mESCs) up- and down-regulated in AHARIBO-rC RNA or global translatome relative to the AHARIBO-nP proteome.

**Figure S4.**
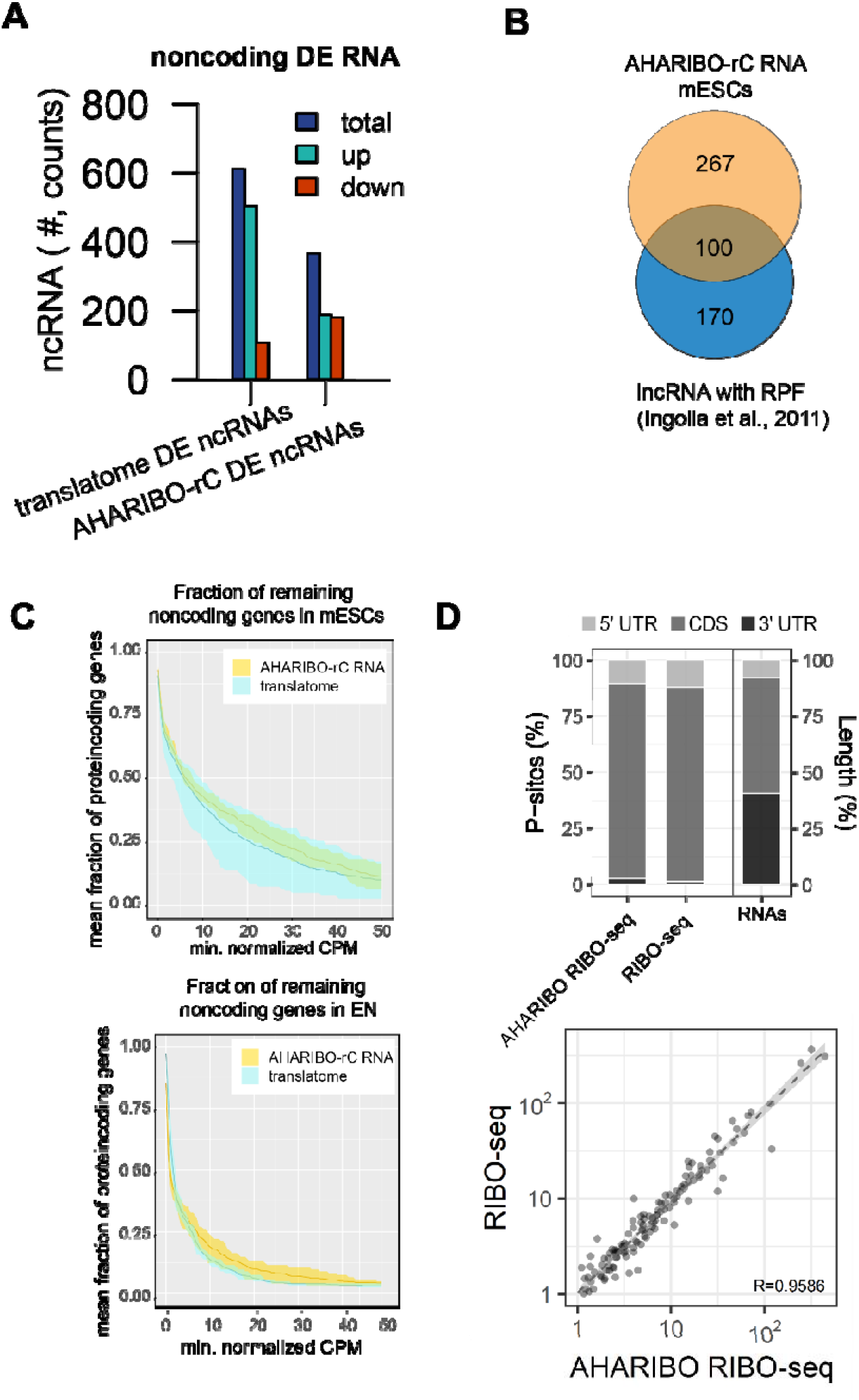
Isolation of lncRNAs with AHARIBO. A) Number of up- and down-regulated differentially expressed non-coding RNAs in the global translatome and in AHARIBO-rC RNA DE, differentially expressed. ncRNA, noncoding RNA. B) Venn diagram representing the number of differentially expressed lncRNAs identified by AHARIBO-rC (orange) and number of lncRNAs with at least 1 CPM in Ingolia et al., 2011 (blue). C) Fraction of remaining non-coding genes at increasing CPM thresholds in AHARIBO-rC versus global translatome RNA from both mESCs (top) and ENs (bottom). The mean of n = 3 biological replicates is reported. Shadows: data are represented as mean ± SD. D) Percentage of ribosome P-sites mapping to the 5’ UTR, coding sequence (CDS), and 3’ UTR of mRNA from AHARIBO RIBO-seq and standard RIBO-seq data. The percentage length of each mRNA region is indicated on the right-hand y axis (top). Data correlation of AHARIBO Ribo-seq and standard RIBO-seq (performed on the input) obtained in mESCs. Results are representative of two independent replicates for each method (bottom).

## Materials and Methods

### Cell culturing and treatments

For protocol development, optimization and validation, HeLa cells were used. HeLa cells were maintained on adherent plates in Dulbecco’s modified Eagle’s medium (DMEM; EuroClone #ECM0728L) supplemented with 10% fetal bovine serum, 2 mM L-glutamine, 100 units/mL penicillin and 100 ug/mL streptomycin at 37°C, 5% CO_2_. For passaging, cells were washed with 1x PBS, detached using 0.25% trypsin-EDTA and spun down at 260 x g for 5 minutes.

For treatments, 250,000-400,000 HeLa cells per well were seeded in 6-well plates and grown to 80% confluence. At the time of treatment, culture medium was removed and cells were washed once with warm 1x PBS. Subsequently, cells were incubated with Dulbecco’s modified Eagle’s limiting medium (DMEM-LM; Thermo Scientific #30030) supplemented with 10% fetal bovine serum and 800 μM L-leucine for 40 min to deplete methionine reserves. Methionine-free medium was then supplemented with L-azidohomoalanine (Click-IT™ AHA; Invitrogen #CI0I02) at a final concentration of 250 μM and incubation time (ranging from 10 min to 120 min; 30 min set as incubation time for the protocol). Cells were then treated with 1x sBlock^®^ (IMMAGINA BioTechnology, catalog n° #RM8; sBlock is an anisomycin-containing proprietary reagent) for 10 min. Then, 6-well plates were placed on ice, medium was removed and cells were washed once with cold 1x PBS supplemented with 1x sBlock. After removing residual PBS with a pipette, hypotonic lysis buffer (0.1M NaCl, 0.1 M MgCl2, 0.1M Tris HCL, 1% Tx-100, 1x sBlock, 1% sodium deoxycholate, 5 units/mL DNAse I – Thermo Scientific #89836, 200 units/mL RiboLock RNase Inhibitor – Thermo Scientific #E00381, 1x Protease Inhibitor Cocktail – Cell Signaling Technology #5871S) was added to each well, and cells were lysed with the aid of a scraper. After hypotonic lysis, nuclei and cellular debris were removed by centrifuging at 18000 x g, 4°C for 5 min. For quantification of the total absorbance value of cell lysates, the absorbance was measured (260 nm) using a Nanodrop ND1000 UV-VIS Spectrophotometer. Lysates were aliquoted and processed directly or stored at −80°C.

Arsenite pre-treatment was performed adding sodium arsenite (Sigma-Aldrich #S7400) at a final concentration of 500 μM for 1 hour.

For RNA-seq and proteomics experiments, two biological settings were assessed in triplicate experiments: i) undifferentiated mouse 46C embryonic stem cells (mESCs) (Ying et al., 2003) and ii) mESCs induced to differentiate into early neurons (ENs).

mESCs were maintained in mESC self-renewal medium composed of Glasgow’s MEM (Thermo Scientific #11710-035) supplemented with 1000 units/ml ESGRO Recombinant Mouse LIF protein (Millipore #ESG1107), 10% fetal bovine serum, 55 μM 2-Mercaptoethanol, 1mM Sodium Pyruvate (Thermo Scientific #11360070), MEM Non-Essential Amino Acids (Thermo Scientific #11140050), GlutaMax (Thermo Scientific #35050061), and penicillin/streptomycin. For passaging, mESCs were washed twice with 1x PBS, detached using 0.02-0.05% trypsin-EDTA and spun down at 260 x g for 3 minutes. Pellet was resuspended in fresh medium and plated onto 0.1% gelatin-coated culture vessels.

For treatments, 5 x 10^5^ mESCs/cm^2^ were seeded in petri dishes and grown to 60% confluence. For pSILAC proteomics, 24 h before lysis mESCs were washed twice with 1x PBS and the medium was replaced with SILAC Advanced DMEM/F-12 Flex Medium (Thermo Scientific #A2494301), supplemented with 1000 units/ml ESGRO Recombinant Mouse LIF protein, 10% dialyzed fetal bovine serum, 4500 mg/L glucose, 17.25 mg/L proline, and penicillin/streptomycin. Either light or heavy L-arginine (Sigma-Aldrich #608033) and L-lysine (Sigma-Aldrich #608041) were added at 84 mg/L and 146 mg/L, respectively. For both AHA+ proteomics and RNA-seq experiments, treatments were performed as described above for HeLa cells, with the exception that methionine-free medium was supplemented with 1000 units/ml ESGRO Recombinant Mouse LIF protein and 10% dialyzed fetal bovine serum. After methionine depletion, cells were treated with 250 μM AHA for 30 min. The remaining treatment steps and hypotonic lysis were performed as detailed above.

Neuronal differentiation was performed according to a previously described protocoling et al., 2003). Briefly, 2.000 mESCs/cm^2^ were seeded on gelatin-coated culture vessels in N2B27 medium. Cells were gently washed with 1x PBS and medium was renewed every 1-2 days until 15DIV. N2B27 medium is composed of 1:1 mix of DMEM/F-12 (Thermo Scientific #21331020) and Neurobasal Medium (Thermo Scientific #21103049), supplemented with 0.5% N-2 (Thermo Scientific #17502048), 1% B-27 (Thermo Scientific #17504044), Glutamax, and penicillin/streptomycin.

Upon differentiation, early neurons were treated directly in culture vessels. For pSILAC proteomics, 24 h before lysis ENs were washed once with 1x PBS and the medium was replaced with SILAC Advanced DMEM/F-12 Flex Medium, supplemented with 0.5% N2, 1% B27, 4500 mg/L glucose, 17.25 mg/L proline, and penicillin/streptomycin, 4500 mg/L glucose, 17.25 mg/L proline, and penicillin/streptomycin. Either light or heavy L-arginine and L-lysine were added at 84 mg/L and 146 mg/L, respectively. For both AHA+ proteomics and RNA-seq experiments, ENs were treated as described above for HeLa cells, with 250 μM AHA for 30 min. The remaining treatment steps and hypotonic lysis were performed as detailed above.

### Immunocytochemistry

For immunofluorescence assay, cells were fixed with 4% paraformaldehyde for 15 min at room temperature, permeabilized using 0.5% Triton X-100 in 1x PBS for 15 min at room temperature and blocked using 5% fetal bovine serum, 0.3% Triton X-100 in 1x PBS for 2h at room temperature. Cultures were then incubated overnight at 4°C with either anti-β3-tubulin (Promega #G712A) or anti-Oct4 (Santa Cruz Biotechnologies #SC-5279) primary antibodies diluted in 2% fetal bovine serum, 0.2% Triton X-100 in 1x PBS. Cells were then washed three times with 1x PBS and incubated with Alexa-555 anti-mouse secondary antibodies for 2h. Nuclei were counterstained with Hoechst 33258 before imaging with a Zeiss Axio Observer Z1 inverted microscope equipped with a 2.83 Megapixel AxioCam 503 mono D camera.

### AHARIBO-rC/AHARIBO-nP: purification of active ribosomes for RNA/protein isolation

For RNAseq experiments, lysates were diluted in W-buffer (10mM NaC1, 10mM MgC_2_, 10 mM Hepes, 1x sBlock) to a final Nanodrop-measured absorbance (260 nm) of 1-2 a.u./mL, supplemented with 40 U of Superase-In RNase Inhibitor (Thermo Scientific #AM2696) and incubated with Dibenzocyclooctyne-PEG4-biotin conjugate (Sigma-Aldrich #760749; 50μM final concentration) in a reaction volume of 100 μl for 1 h on a rotator in slow motion (9 rpm) at 4°C. Lysates were then incubated with 50 μl of eMagSi-cN beads (IMMAGINA BioTechnology #018-eMS-001) for 30 min at 4°C on the rotator in slow motion (9 rpm). Subsequently, samples were taken off the rotator and placed on a magnetic rack on ice, and supernatants were discarded. Beads were washed two times with 500 μl of 1x PBS supplemented with 0.1% Triton-X100, 1x sBlock and 1:10,000 RiboLock RNase Inhibitor (Thermo Scientific #E00381) on the rotator in slow motion at 4°C, removing supernatants from the tubes sitting on the magnetic rack and gently adding new washing solution each time. After the final wash, beads were resuspended in 200 μl of W-buffer and transferred to a new vial. Then, 20 μl of 10% SDS and 5 ul of Proteinase K (Qiagen #19131) were added to each sample, and samples were incubated at 37°C for 75 min in a water bath. Subsequently, suspensions were transferred to a new vial and acid phenol:chloroform:isoamyl alcohol RNA extraction was performed. Briefly, an equal volume of acid phenol:chloroform:isoamyl alcohol (pH 4.5) was added, and samples were vortexed and centrifuged at 14,000 x g for 5 min. Aqueous phases were then transferred to new vials, 500 μl of isopropanol and 2 μl of GlycoBlue (Thermo Scientific #AM9516) were added, samples were mixed and incubated at room temperature for 3 min and then stored overnight at – 80 °C. The following day, samples were centrifuged at 14,000 x g for 30 min, supernatants were removed, 500 μl of 70 % ethanol were added to each sample and samples were then centrifuged at 14,000 x g for 10 min. Finally, pellets were air-dried and resuspended in 10 μl of nuclease-free water. When quality check and quantification was needed, RNA samples were run on a 2100 Bioanalyzer (Agilent) using the Agilent RNA 6000 Nano Reagents kit (Agilent #5067-1511) and assayed on the Qubit fluorometer using the Qubit RNA HS Assay Kit (Thermo Scientific # Q32852). For visualization of total RNA patterns, samples were run on a 1% agarose gel. ImageJ software (v 1.45s) was used for quantitation of signal intensities of ribosomal RNA bands.

For proteomics experiments, lysates were diluted in W-buffer to a final Nanodrop-measured absorbance (260 nm) of 1-2 a.u./mL in a final volume of 100 ul. Ribosome pulldown was performed using Mag-DBCO beads (IMMAGINA BioTechnology #MDBCO). Lysates were incubated with 50 μl of beads for 1h on a rotator in slow motion (9 rpm) at 4°C. Supernatants were discarded after placing samples on the magnetic rack. Beads were washed three times with 500 μl of 200 mM Tris, 4 % CHAPS, 1 M NaCl, 8M Urea, pH 8.0 at room temperature on a shaker at 1000 rpm, using the magnetic rack to replace the washing solution. After the final wash, beads were resuspended in 30 μl of water and transferred to a new vial.

### qRT-PCR analysis

Total RNA was extracted from samples processed through the AHARIBO-rC protocol as described above. Depending on the available input material, RNA was retrotranscribed using either RevertAid First Strand cDNA Synthesis Kit (Thermo Scientific #KI621) or Superscript™ III Reverse Transcriptase (Thermo Fisher #18080044), per manufacturer protocols. qPCR was run on CFX Connect Real-Time PCR Detection System (BioRad) using Kapa Probe Fast Universal qPCR Kit (Kapa Biosystems #KK4702). Reactions were performed in technical duplicates of biological triplicates. The following TaqMan probes were used: Hs99999901_s1 (18S), Hs02800695_ml (HPRT1).

For normalization of qRT-PCR results, HPRT1 was used as housekeeping gene. The fold change in normalized 18S mRNA levels between untreated (control) and treated (arsenite) samples was calculated. A second normalization to threshold cycles from non-AHA-treated samples was done to account for background signal.

### RNA-seq

RNA samples were subjected to library preparation for the Illumina platform using the SMART-Seq Stranded Kit (Takara #634443) as manufacturer instructions, using 5 ng of RNA as starting material. For quality check and quantification, the final libraries were run on a 2100 Bioanalyzer (Agilent) using the Agilent DNA 1000 Kit (Agilent #5067-1504) and assayed on the Qubit fluorometer using the Qubit dsDNA HS Assay Kit (Thermo Scientific # Q32851). Libraries were sequenced on an Illumina HiSeq2500 by the NGS Core Facility (University of Trento).

### Polysome profiling

HeLa cells were treated and lysed as described above, adding one the following blocking drugs: i) sBlock (IMMAGINA BioTechnology #SM8, final concentration 1x, 10 min treatment); ii) cycloheximide (Sigma-Aldrich #C4859; final concentration 36 μM, 5 min treatment); iii) puromycin (Sigma-Aldrich #P8833; final concentration 50 μM, 5 min treatment); iv) no blocking drug. Cleared supernatants obtained from cytoplasmic lysates were loaded on a linear 15%-50% sucrose gradient and ultracentrifuged in a SW41Ti rotor (Beckman) for 1 h and 40 min at 180,000 x g at 4°C in a Beckman Optima LE-80K Ultracentrifuge. After ultracentrifugation, gradients were fractionated in 1 mL volume fractions with continuous absorbance monitoring at 254 nm using an ISCO UA-6 UV detector. Each fraction was flash-frozen in liquid nitrogen and stored at – 80°C for subsequent protein extraction.

Polysome profiles were analyzed as follows. The relative intensity of each individual fraction was determined for both on-membrane AHA and RPL26 signals, then the AHA/RPL26 relative intensity ratio was calculated for each fraction. For each profile, the relative intensity ratios of polysome-containing fractions (fractions 8/9-10/11) were averaged and normalized to the relative intensity ratio of the 60S fraction, which was chosen as internal baseline for background signal based on the fact that it should be devoid of translationally active ribosomes. To assess the effect of the different blocking drugs, averaged normalized relative intensity ratios for the profiles obtained from different blocking drugs and from the untreated control sample were compared. ImageJ software (v 1.45s) was used for quantitation of signal intensities of protein bands.

### Sucrose cushioning for ribosome enrichment (global translatome)

HeLa cells were treated in petri dishes and lysed as described above, adding 1x sBlock as blocking drug. Sucrose cushioning was performed according to a modified version of a previously described protocol (Ingolia et al., 2012b). For each sample, a volume of cell lysate corresponding to 1.7 a.u. (based on Nanodrop measurement of absorbance at 260 nm) was layered on top of 900 μl of 30% sucrose cushion (30 mM Tris-HCl pH 7.5, 100 mM NaCl, 10 mM MgCl_2_, 1M sucrose in nuclease-free water) supplemented with 1x sBlock. Samples were ultracentrifuged at 95,000 rpm at 4°C for 1 h and 40 min using a TLA100.2 rotor (Beckman). Supernatants were saved if needed for further analysis, and pellets were resuspended in 100 μl of nuclease-free water supplemented with 30 mM Tris-HCl pH 7.5, 100 mM NaCl, 10 mM MgCI_2_.

### Protein extraction from sucrose gradient fractions

Polysomal fractions (1 mL) or pellet/supernatant fractions from 30% sucrose cushioning (1/5^th^ of total amount, adjusted to 260 μl volume) were processed for methanol/chloroform protein extraction. Briefly, 600 μl of methanol and 150 μl of chloroform were added to each sample and samples were vortexed. Then, 450 μl of deionized water were added to each sample and samples were vortexed again. Samples were centrifuged at 14,000 x g for 1 min at room temperature and the resulting aqueous phase was removed without disrupting the underlying white ring (protein interface). Subsequently, 450 μl of methanol were added to each sample, samples were vortexed and then centrifuged at 14,000 x g for 2 min at room temperature. After centrifugation, supernatants were removed and pellets air-dried. Finally, pellets were resuspended in deionized water supplemented with Pierce Lane Marker Reducing Sample Buffer (Thermo Scientific #39000) to a final volume of 15 μl and either stored at −80°C or heated at 95°C and directly used for SDS-PAGE.

### On-membrane click chemistry

Cell lysate or protein extracts obtained from sucrose gradient fractions were supplemented with Pierce Lane Marker Reducing Sample Buffer (Thermo Scientific #39000), heated at 95°C for 10 min and separated by SDS-PAGE. Separated proteins were transferred to nitrocellulose membranes, then membranes were blocked overnight at 4°C in 5% milk prepared in 1x TBS – 0.1% Tween20 supplemented with Dibenzocyclooctyne-PEG4-biotin conjugate (Sigma-Aldrich #760749; 50μM final concentration). Membranes were washed three times in 1x TBS – 0.1% Tween20 for 10 min each, then incubated with Precision Protein StrepTactin-HRP Conjugate (BioRad #1610380; 1:1000 in 5% milk prepared in 1x TBS – 0.1% Tween20) for 1 h at room temperature, then washed again. Membranes were subsequently developed using Amersham ECL Prime Western Blotting Detection Reagent (GE Healthcare #RPN2236). Images were acquired through the ChemiDoc MP Imaging System. ImageJ software (v 1.45s) was used for quantitation of AHA signal intensities.

### Immunoblotting

10-20 μl of cell lysate or protein extracts obtained from sucrose gradient fractions were supplemented with Pierce Lane Marker Reducing Sample Buffer (Thermo Scientific #39000), heated at 95°C for 10 min and separated by SDS-PAGE. Separated proteins were transferred to nitrocellulose membranes, then membranes were blocked for 1 h at room temperature in 5% milk prepared in 1x TBS – 0.1% Tween20. Membranes were subsequently incubated for 1 h at room temperature with the following primary antibodies, diluted in 5% milk prepared in 1X TBS – 0.1% Tween20: anti-RPL26 (Abcam #ab59567; 1:2000), anti-RPS6 (Abcam #ab40820; 1:1000), anti-eEF2 (Abcam #ab33523; 1:1000), anti-beta-actin (Abcam #ab8227; 1:2000). Membranes were washed three times in 1x TBS – 0.1% Tween20 for 10 min each, then incubated with the appropriate HRP-conjugated secondary antibodies for 1h at room temperature and washed again as before. Membranes were then developed using either Amersham ECL Prime Western Blotting Detection Reagent (GE Healthcare #RPN2236) or SuperSignal West Femto Maximum Sensitivity Substrate (Thermo Scientific #34095), depending on signal intensities. Images were acquired through the ChemiDoc MP Imaging System. ImageJ software (v 1.45s) was used for quantitation of signal intensities of protein bands.

### RNA-seq data analysis

FASTQ files from Illumina HiSeq2500 were firstly checked for adapters and quality-base distribution using FASTQC tool (http://www.bioinformatics.babraham.ac.uk/projects/fastqc) followed by trimming with Trimmomatic-0.35(Bolger et al., 2014) with the following settings: ILLUMINACLIP:ADAPTOR_FILE:2:30:8:1 LEADING:3 TRAILING:3 SLIDINGWINDOW:4:15. Prior to sequencing data processing, technical replicates (different sequencing lanes) from the same library were merged together generating a unique FASTQ per sample. Reads were aligned onto mm10 Mouse genome using STAR-2.6.0a (Dobin et al., 2013) with a maximum mismatch of two and default setting for all other parameters. Once uniquely mapped reads were selected, the GRCm38.92 mouse gene annotation from Ensembl (www.ensembl.org) was incorporated in the HTSeq-count v0.5.4 (Anders et al., 2015) tool to obtain gene-level counts. Genes with CPM (Counts Per Million) < 1 in all samples were considered as not expressed and hence removed from the analysis. TMM (Trimmed Mean of M values) (Robinson and Oshlack, 2010) normalization and CPM conversion were then performed to obtain gene expression levels for downstream analyses. For each comparison, differential expression testing was performed using the edgeR-3.20.9 (Robinson et al., 2010) statistical package from Bioconductor. According to the edgeR approach, both common (all genes in all samples) and separate (gene-wise) dispersions were estimated and integrated into a Negative Binomial generalized linear model to moderate gene variability across samples. Finally, genes having a Log fold-change higher/smaller than 1.5/-1.5 and a FDR-corrected P-value of 0.01 (or smaller) were considered as significant.

### Proteomics experiments

Proteomic analysis was performed on samples processed through the pSILAC and AHARIBO workflows, as described above. For pSILAC experiments, cells were prepared as described above (see Cell culturing and treatments). 50□μg of lysates were then subjected to acetone precipitation and protein pellets were dissolved in 50 mM ammonium bicarbonate and 6M urea. For AHARIBO enrichment, the beads used for ribosome pulldown were reconstituted in 100 μl 50 mM ammonium bicarbonate.

Samples were reduced using 10 mM DTT for 1 h at room temperature and alkylated with 20 mM iodoacetamide in the dark for 30 min at room temperature. Subsequently, proteins were digested at room temperature with 0.5 μg Lys-C (Promega, # VA1170) for 4 h, after which the solution was diluted 4 times in 50 mM ammonium bicarbonate. 1 μg of trypsin (Promega, # V5111) was then added to the samples and proteolysis was carried out overnight. Digestion was interrupted by adding 1% trifluoroacetic acid. Samples were then desalted by C18 stage-tip, lyophilized, and resuspended in 20 μl of buffer A (0.1% formic acid) for LC-MS/MS analysis.

Samples were analyzed using an Easy-nLC 1200 system coupled online with an Orbitrap Fusion Tribrid mass spectrometer (both Thermo Fisher Scientific). Peptides were loaded onto a 25 cm long Acclaim PepMap RSLC C18 column (Thermo Fisher Scientific, 2μm particle size, 100Å pore size, id 75 μm) heated at 40°C. For pSILAC samples, the gradient for peptide elution was set as follow: 5- to 25% buffer-B (80% acetonitrile, 0.1% formic acid) over 90 min, 25 to 40% over 15 min, 40% to 100% over 18 min and 100% for 17 min at a flow rate of 400 nl/min. For AHARIBO pulldown samples, the same steps for peptide elution were set over a total gradient of 80 min. The instrument was set in a data-dependent acquisition mode. The full MS scan was 350-1100 m/z in the orbitrap with a resolution of 120,000 (200 m/z) and an AGC target of 1×10e6. MS/MS was performed in the ion trap using the top speed mode (3 secs), an AGC target of 5×10e3 and an HCD collision energy of 30.

MS raw files were analyzed by using Proteome Discoverer (v2.2, Thermo Scientific). MS/MS spectra were searched by the SEQUEST HT search engine against the human or the mouse UniProt FASTA databases (UniProtKB 11/2018). Trypsin was specified as the digestive enzyme. Cysteine carbamidomethylation (+57.021 Da) was set as fixed modification, methionine oxidation (+15.995 Da) and N-term acetylation (+42.011 Da) as variable modifications. SILAC labeling (Lys +8.014 Da, Arg +10.008 Da) was used as quantification method for pSILAC samples. All other values were kept as default.

### Proteomics data analysis

Heteroscedastic T-test was used to assess the significant differences in peptide/protein abundance (p-value lower than 0.05) unless stated otherwise. Data distribution was assumed to be normal but this was not formally tested. Gene Ontology (GO) and KEGG (Kyoto Encyclopedia of Genes and Genomes) pathway analysis were performed using DAVID version 6.8 and PANTHER 14.1 and Enrichr (http://amp.pharm.mssm.edu/Enrichr/).

### Identification of IncRNA peptides from result spectra

Sequenced non-coding RNAs were in-silico translated into amino acid sequences using the EMBOSS Transeq tool from EMBL. Only the three forward frames were translated. Spectra obtained from the AHARIBO enrichment of newly synthesized proteins were searched against a database of typical contaminants like keratins, trypsin, bovine serum albumin etc provided by MaxQuant (PMID: 19029910). The software utilized for database searching was Proteome Discoverer (v2.4, Thermo Scientific); the Non-fragment filter and the Top N Peaks Filter (with N=4 per 100 Da) were also used in the workflow to eliminate noise signals from the MS/MS spectra. The spectra not matching with high confidence this database were searched against the human SwissProt database. Those not matching with both databases were used to match the in-silico translated database generated by EMBOSS Transeq using semi-specific tryptic cleavage to consider also unexpected translation start sites. We considered only those spectra that passed the 1% FDR threshold, and we created two distinct groups for those peptides with an AUG “in-frame” vs not in-frame.

### Ribosome Profiling

mESCs at 80% confluence were pre-treated with the elongation inhibitor cycloheximide before rapid harvest on ice and cell lysis (lysis buffer, IMMAGINA BioTechnology #RL001-1). Clarified cell lysates (1.7 total a.u., measured by Nanodrop) were treated with 1.3 U of RNase I (Thermo, #AM2295) in W-buffer (IMMAGINA BioTechnology #RL001-4) containing 1x sBlock to digest RNA not protected by ribosomes. Digestion was performed for 45 min at RT and then stopped with Superase-In RNase Inhibitor (Thermo Scientific #AM2696) for 10 min on ice. Samples were then processed differentially according to the specific approachas described below.

#### Standard RIBO-seq

80S ribosomes were isolated by centrifuging lysates through a 30% sucrose cushion at 95000 rpm, for 2 h at 4°C. The cushion was resuspended in W-buffer and treated with SDS 10% (final 1%) and 5 μL of proteinase K (20mg/mL), and incubated at 37 °C in a waterbath for 75 min before acid phenol:chlorophorm:isoamyl alcohol (pH 4.5) RNA extraction.

#### AHARIBO RIBO-seq

The mixture was incubated with Dibenzocyclooctyne-PEG4-biotin conjugate (Sigma-Aldrich #760749; 50 μM final concentration) in a reaction volume of 100 μl for 1 h on a rotator in slow motion (9 rpm) at 4°C. Lysates were then incubated with 50 μl of eMagSi-cN beads (IMMAGINA BioTechnology #018-eMS-001) for 30 min at 4°C on the rotator in slow motion (9 rpm). Subsequently, samples were taken off the rotator and placed on a magnetic rack on ice, and supernatants were discarded. Beads were washed two times with 500 μl of 1x PBS supplemented with 0.1% Triton-X100, 1x sBlock and 1:10,000 RiboLock RNase Inhibitor (Thermo Scientific #E00381) on the rotator in slow motion at 4°C, removing supernatants from the tubes sitting on the magnetic rack and gently adding new washing solution each time. After the final wash, beads were resuspended in 200 μl of W-buffer and transferred to a new vial. Then, 20 μl of 10% SDS and 5 ul of Proteinase K (Qiagen #19131) were added to each sample, and samples were incubated at 37°C for 75 min in a water bath. Subsequently, suspensions were transferred to a new vial and acid phenol:chloroform:isoamyl alcohol (pH 4.5) RNA extraction was performed.

For both approaches, protocol steps starting from phenol:chloroform:isoamyl alcohol RNA extraction were performed as follows. Briefly, an equal volume of phenol:chloroform:isoamyl alcohol was added, and samples were vortexed and centrifuged at 14,000 x g for 5 min. Aqueous phases were then transferred to new vials, 500 μl of isopropanol and 2 μl of GlycoBlue (Thermo Scientific #AM9516) were added, samples were mixed and incubated at room temperature for 3 min and then stored overnight at – 80 °C. The following day, samples were centrifuged at 14,000 x g for 30 min, supernatants were removed, 500 μl of 70 % ethanol were added to each sample and samples were then centrifuged at 14,000 x g for 10 min. Finally, pellets were air-dried and resuspended in 10 μl of nuclease-free water. Extracted RNA was then resolved by electrophoresis through denaturing TBE-urea gels, and fragments between 25 nt and 35 nt were excised. Libraries were prepared using the RiboLace kit_module 2 (IMMAGINA BioTechnolgy #RLOOl_mod2) and sequenced on an Illumina HiSeq 4000 sequencer with a single-end 50 base pair run.

### RIBO-seq data analysis

Reads were processed by removing 5□ adapters, discarding reads shorter than 20 nucleotides and trimming the first nucleotide of the remaining ones (using Trimmomatic v0.36). Reads mapping on the collection of M. musculus rRNAs (from the SILVA rRNA database, release 119) and tRNAs (from the Genomic tRNA database: gtrnadb.ucsc.edu/) were removed. Remaining reads were mapped on the mouse transcriptome (using the Gencode M17 transcript annotations). Antisense and duplicate reads were removed. All alignments were performed with STAR (v020201) employing default settings.

The identification of the P-site position within the reads was performed using riboWaltz (v1.1.0) computing the P-site offsets from the 3’ end of the reads. The percentage of P-sites falling in the three annotated transcript regions (5’ UTR, CDS and 3’ UTR) was computed using the function *region_psite* included in riboWaltz (v1.1.0).Transcript counts were normalized using the trimmed mean of M-values normalization method (TMM) implemented in the edgeR Bioconductor package. Transcripts displaying 1 CPM in at least one replicate were kept for further analyses.

## ADDITIONAL INFORMATION

### Author contributions

L.M. suggested the original workflow. M.C. conceived the experiments. A.P. prepared the RNA libraries for RNA-seq. C.F, L.M, A.P., A.D.P performed the experiments. R.B, D.P, performed proteomics experiments. J.Z., cultured mESC cells, performed the 10-day neuronal differentiation and collected immunofluorescence images. A.Q suggested the biological model of neuronal differentiation and made available mESC cells. M.C., S.S, R.B, D.P analyzed proteomic data. SS identified lncRNA peptides from result spectra. F.G, A.R, M.C. analyzed RNA-seq data. G.V. and F.L. analyzed RIBO-seq and AHARIBO RIBO-seq data. L.M., A.P., M.C. drafted the manuscript. MC and GV reviewed the manuscript. M.C., A.P, P.B., L.M., S.S, A.R., C.F., A.D.P., J.Z., F.G, G.V, G.G. reviewed the final manuscript. All of the authors read and approved the final manuscript.

