## Supplementary figures and images for "One-shot analysis of translated mammalian lncRNAs with AHARIBO"

### S1.tif

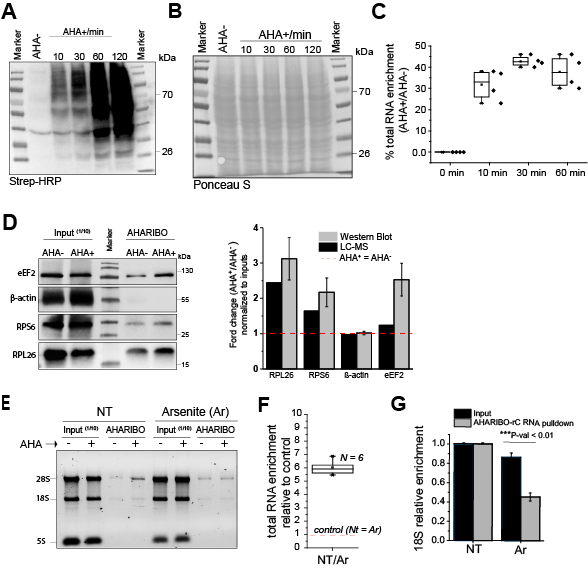

### S2.tif

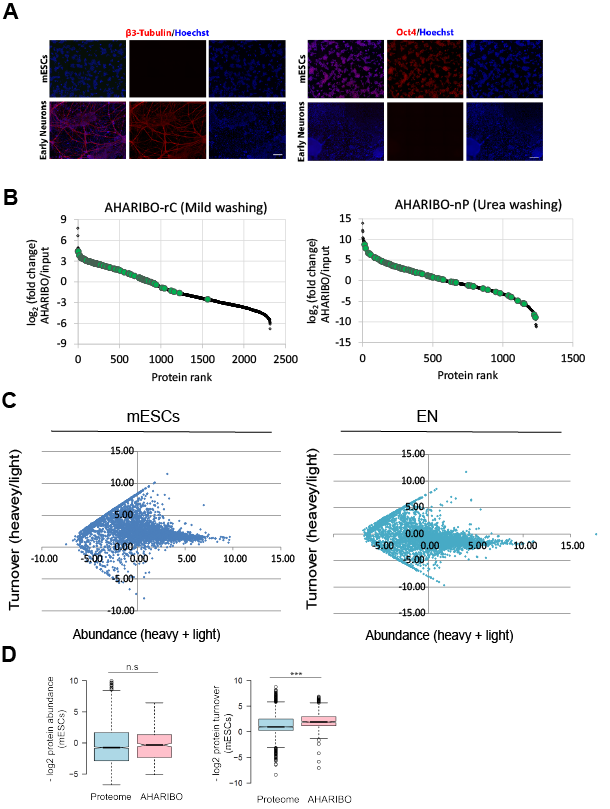

### S3.tif

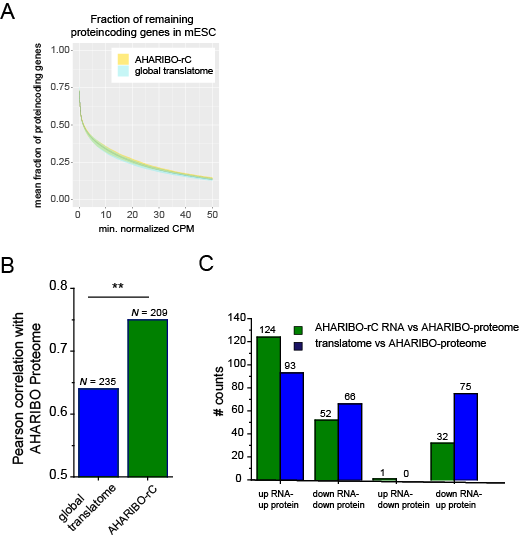

### S4.tif

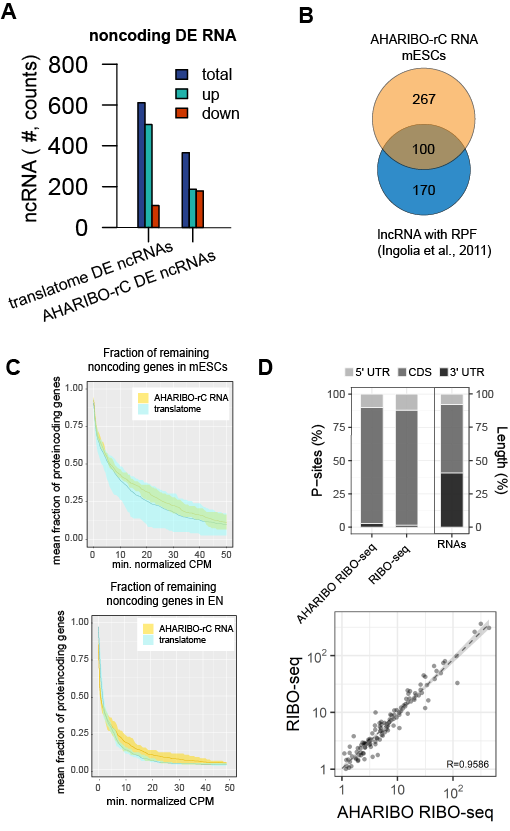
